# Gene-centric constraint of metabolic models

**DOI:** 10.1101/116558

**Authors:** Nick Fyson, Min Kyung Kim, Desmond S. Lun, Caroline Colijn

## Abstract

**Motivation:** A number of approaches have been introduced in recent years allowing gene expression data to be integrated into the standard flux Balance Analysis (FBA) technique. This additional information permits greater accuracy in the prediction of intracellular fluxes, even when knowledge of the growth medium and biomass composition is incomplete, and allows exploration of organisms’ metabolism under wide-ranging conditions. However, existing techniques still focus on the reaction as the fundamental unit of their modelling. This carries the advantages of incorporating expression measurements, but discounts the fact that genes (and their associated proteins) may be involved in the catalysis of multiple reactions through the formation of alternative protein complexes.

**Results:** We demonstrate an approach focusing not on reactions or genes as the fundamental unit, but on the ‘Gene Complex’ (GC), a set of genes that is sufficient to catalyse a given reaction. We define expression-based limits in such a way that proteins cannot do ‘double duty’: no single molecule is permitted to contribute to the catalysis of more than one reaction at a time. Using experimentally determined RNA expression and intracellular fluxes, we validate this novel and more conceptually sound approach.

**Availability and Implementation:** An implementation of the GC-Flux algorithm is available as part of the Pyabolism python module. https://github.com/nickfyson/pyabolism

## 1 Introduction

Modelling the network of metabolic reactions inside cells is key to understanding their behaviour, and recent years have seen a diverse range of approaches tackling the problem [7]. Fundamentally, the metabolism of a cell can be viewed as a large system of differential equations, but limitations on availability and accuracy of the necessary data means such full-cell dynamic models remain rare [27]. Consequently, we have seen widespread development of Constraint-Based Models (CBMs), based on the foundation of flux Balance Analysis (FBA) [41]. FBA does not require knowledge of reaction kinetics, instead relying on the evolutionary assumption that cells will exhibit near-optimal behaviour under their internal and externally imposed constraints. Crucially, this requires only the list of all reactions present inside a given cell, information for which acquisition may be largely automated.

Constraint-based modelling is well established [43, 45, 41]; the framework is as follows. Given a comprehensive set of all reactions within a cell, R, the state of the cell's metabolism can be represented by a vector of fluxes, one for each reaction. In isolation the reactions are free to run at any thermodynamically feasible rate, but if we assume that the cell is operating at steady state, with no overall changes in the concentrations of the metabolites, this imposes a large set of linked constraints on the flux vector — the fluxes must be such that the total of each metabolite generated is equal to that consumed. This network of reactions can be represented by the stoichiometric matrix, *S*, in which each row represents a metabolite and each column gives the stoichiometry for a particular reaction.

To simulate the growth of cells, some additional special reactions are needed. ‘Exchange reactions’ represent the input/output of metabolites from/to the outside world, where the metabolite is only present on one side of the reaction. These permit an overall imbalance in specific metabolites — such as the primary carbon source — and can be constrained to represent particular growth conditions. Finally, the standard FBA approach requires a ‘biomass reaction’ which details the range and precise quantity of metabolites that go into constructing a new gram of cellular biomass.

A Linear Program (LP) is then constructed using the stoichiometry and exchange bounds as constraints, and the flux through the biomass vector as the objective function. We follow the standard notation for FBA modelling, where *S* the stoichiometric matrix and *v* the vector of reaction fluxes. The objective for optimization is defined by a vector *c*, which gives a weighting to each of the reaction fluxes, and in the standard FBA formulation will have a single non-zero entry corresponding to the biomass reaction. Finally each reaction *i* has a lower bound *a_i_* and an upper bound *b_i_*. The optimisation problem can then be defined as

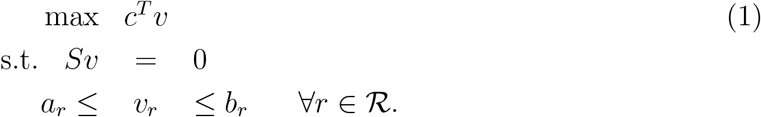

This basic formulation has been widely used for modeling bacterial growth for decades (e.g. [41, 45, 15, 3, 51, 7] and many others). In early studies the flux space was constrained by limiting only the uptake of nutrients [14, 48, 6]. Subsequent studies showed that it is possible to make quantitative predictions of internal flux distributions using CBMs, both with more sophisticated use of the basic stoichiometric model [34, 50, 45], and through combination with increasingly abundant high-throughput expression data (e.g. [10, 34, 52] and many others).

Transcriptomic data define an in-depth, quantitative and genome-scale description of cellular activity, and as a result, integrating transcriptomic measurements with constraint-based modeling (CBM) in principle allows researchers to compare an organism's metabolic function in a wide range of different states. For this reason, there has been considerable interest in integrating ‘omics data with constraint-based metabolic modeling [47, 23, 43, 56], and numerous methods for transcriptomic data are available [2, 31, 30, 12, 4, 17, 10, 52, 57, 25, 9, 47, 23, 44, 5, 11]. One study has examined a wide range of different approaches for integrating expression data with constraint-based modelling, and found no consistent performance benefit, in terms of predicting measured internal fluxes, over the more basic FBA approach [35]. However, the integration of transcriptomic data allows interrogation of situations that cannot be addressed by pFBA ([34]) or similar methods, because these methods do not capture changes in internal regulation that drive differences in metabolism.

There have been many successful applications of tools incorporating transcriptomic data with CBM, despite the fact that mRNA measurements do not directly reflect either protein abundances or, typically, measured flux values (though there are some correlations). [37] used an integrated approach to characterize phenotypes of *Yersinia pestis* under temperature changes which induce virulence and under antibiotic stress. [53] predicted starch content through time in *Ostreococcus tauri,* a model alga, and validated their predictions experimentally; the main regulatory targets in the starch production pathway were correctly predicted. [16] characterized the metabolic adaptations of *M. tuberculosis* to hypoxia. [20] found 88 altered reaction fluxes that characterized a sub-optimal growth state of the cellulolytic anaerobe *C thermocellum.* [1] used metabolic modeling to study the switch from biomass optimization to antibiotic production in *Streptomyces coelicolor,* and found that predicted fluxes showed high correlations with gene expression data. [40] compared two related lineages of *P. aeruginosa* and explored how bacterial metabolism changes over time in cystic fibrosis lungs. [32] examined how *Phaeodactylum tricornutum* increases lipid production, not by a change in growth medium or removal of a reaction but by a remodeling of internal constraints. Transcriptomic data are also incorporated in tools for strain design and engineering [28].

One remaining challenge in integrating transcriptomic (expression) measurements with constraint-based models, despite the successes to date, is that the mapping between genes whose mRNA is measured and reactions in the CBM is not one-to-one. In this paper we introduce a novel approach, introducing the concept of a Gene Complex (GC) and showing how such a shift in focus can improve the prediction of internal fluxes in the hardest of prediction tasks.

### 1.1 E-Flux and E-Flux2

The E-Flux algorithm of Colijn et al. is an extension to basic FBA methodology, permitting the integration of gene expression data into modelling of cellular metabolism without first converting expression into Boolean on/off values [10]. The basic idea relies on the fact that reactions inside cells are catalysed by enzymes, and these enzymes consist of proteins, which are in turn generated through the expression of RNA. The exact quantitative relationship is complicated and generally unknown, but higher intracellular concentration of a given RNA molecule will to some extent be associated with a higher concentration of the corresponding enzyme [19, 18, 38, 39]. This in turn will lead to a greater potential capacity for the reaction, since there is more of the necessary catalyst available.

E-Flux therefore takes experimentally determined data on RNA expression for a particular growth state of a cell, and uses this to set maximum capacities on the *internal* reactions in the constraint-based model. The core E-Flux method chooses the maximum flux, *b_r_*, for the *r*th reaction according to a function of the expression of gene *r* and associated genes

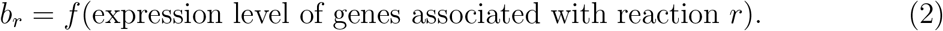

If the reaction catalyzed by the corresponding enzyme is reversible then *a_r_* = −*b_r_*, otherwise *a_r_* = 0. For the standard E-Flux approach, ‘associated genes’ refers both to genes that are components of the same enzyme complex, and genes associated with separate isozymes of the reaction. Note that this is not the same as assuming that expression levels determine realised fluxes, only maximal flux capacities.

The Gene-Protein-Reaction (GPR) associations for a metabolic model are normally coded using a field in the notes of a reaction labelled ‘GENE_ASSOCIATION’, which while not (yet) part of the standard SBML specification has been widely adopted due to implementation in the popular COBRA toolbox. A typical GPR string found in the standard *E. coli* model [42] is

`(( b0978 and b0979 ) or ( b0733 and b0734 )).`

This indicates that the reaction requires one of two enzyme complexes in order to run, either the complex consisting of **b0978** and **b0979**, or the complex consisting of **b0733** and **b0734**. The E-Flux approach uses the expression of RNA as a proxy for the availability of the coded protein, which gives a single number associated with each gene. To translate these into a bound for a reaction with the GPR shown above the following expression is used.

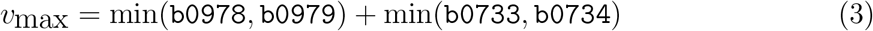

The ‘**and**’ element of the GPR string become min expressions, since the total quantity of the enzyme complex cannot be higher than any one of the constituent proteins. The ‘**or**’ becomes ‘+’ in the expression, since both enzyme complexes can contribute individually to the maximum rate of the reaction. Using this approach we can determine upper bounds on the reaction rate for all reactions in the model that have appropriate transcriptomic data, and then optimise for biomass Flux to find the maximum permitted growth rate.

When looking to predict a specific flux distribution, however, we need to take further steps to ensure a unique distribution. In general there will be a space of fluxes that correspond to the maximal objective value, and the recent paper published by Kim et al. outlines an extension to the original E-Flux algorithm [29]. The E-Flux2 approach consists first of creating a model in which all a priori information on cellular uptake rates and ATP maintenance flux have been removed, before performing a two-stage optimization. The the achieved objective flux from standard E-Flux is added as an additional constraint to the LP, and then the the Euclidean (or L2) norm of the flux vector is minimized. This returns a unique solution, and [29] demonstrated that the resulting flux distribution was significantly correlated with that observed in experiment.

### 1.2 Limitations of E-Flux2

The problem of setting principled bounds for the reactions is complicated when genes appear more than once in the GPR expressions.

(`( b2762 and b2582 ) or ( b2762 and b3781 )).`

In this case we should not simply replace the **or** statement with an addition, since the gene **b2762** appears in both clauses. For this reaction we could solve the problem by nesting our min functions, but there are further cases in which genes appear not just multiple times in a single GPR string, but across multiple reactions within the model.

The E-Flux2 approach has been shown to allow prediction of internal fluxes [29], but it treats all the GPR associations in isolation, permitting enzymes to do ‘double duty’. It is effectively assumed that one protein can be taking part in multiple enzyme complexes, catalysing more than a single reaction at a time. To resolve this issue we restructure the problem, addressing the metabolite network not in a reaction-centric manner, but instead focusing on the Gene Complex (GC).

## 2 Methods

The primary contribution of this paper is the GC-Flux algorithm, but both this novel approach and the existing E-Flux require processing of GPR strings in their implementation. We therefore discuss how the established DAG representation of logical expressions is related to the implementation of our algorithms.

### 2.1 Tree Representation of GPR Strings

While the format for GPR strings found in the Cobra toolbox has seen wide adoption, this still leads to a wide range of equivalent ways to encode relations. Brackets are used to group terms in the GPR string, and hence parsing the string correctly takes some care. Parsing logical expressions is a recognised problem in computer science, and the standard approach is to convert the string into a Directed Acyclic Graph (DAG). In Fig. 1 we give an example of this established approach, showing how a GPR string with brackets may be converted into a DAG representation (in this case a tree), with each gene represented as a leaf node. Internal nodes are labelled by the logical operation associated with that clause in the string, either ‘**OR**’ or ‘**AND**’. In cases where both logical operators appear in a single clause convention dictates that **AND** take precidence, but in practice published metabolic models tend to use brackets to avoid any potential confusion.

**Figure 1:**
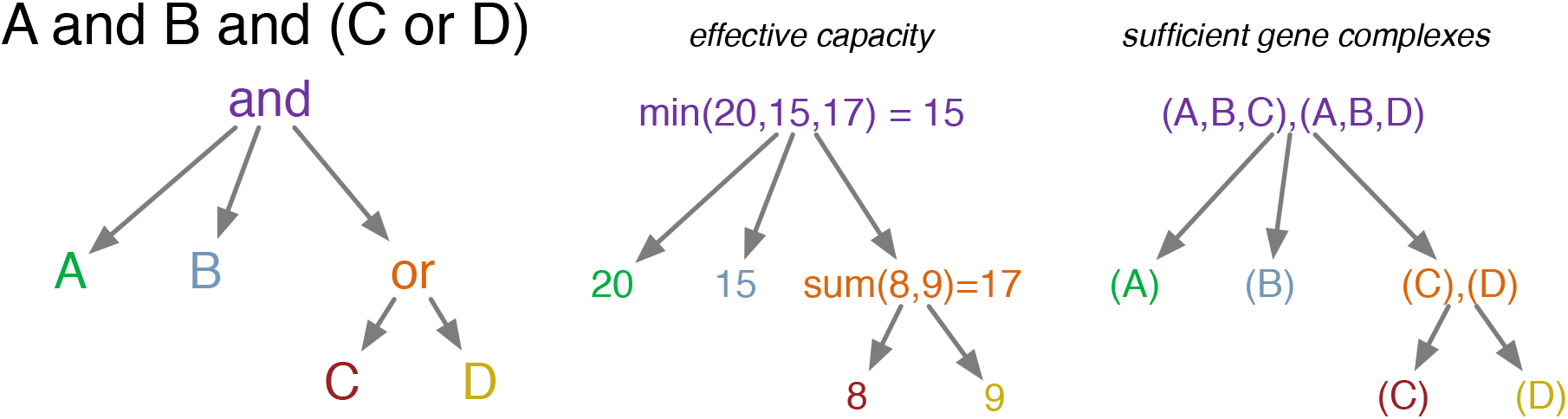
Alternative representation of GPR information as a Directed Acyclic Graph (DAG). While the information is the same, converting into a DAG representation makes it easier to determine effective capacity and viable gene-complexes for a given string.

Representing the GPR string as a DAG clarifies the situation when determining key properties for implementing E-Flux and GC-Flux. For E-Flux we need to determine the maximum quantity of viable catalyst for the reaction, which will guide how fast the reaction is allowed to run. In the DAG representation each unique gene in the GPR string corresponds to a leaf node, and we can add the expression data as a property of each leaf. All the internal nodes are either **AND** or **OR** nodes, and determine how the expression of its child should be calculated. For **AND** nodes, expression is the minimum across all children. For **OR** nodes the effective value of expression is the sum across children. Iterating over the network from leaf to root in reverse topological order will thus allow us to calculate the effective capacity for a reaction as afforded by its GPR string and expression data.

In implementing the GC-Flux algorithm we face a slightly different problem, in which it is necessary to determine a list of gene complexes that are each individually sufficient for catalysis. The full list of viable gene-complexes can be built up from leaves to root, following rules for combining the children of internal nodes. In Fig. 1 we see an example of the approach, with **OR** nodes with two leaf children resulting in a list of two separate complexes, while **AND** nodes lead to each complex being enlarged (while the total number remains the same). An example implementation with full details can be found in the Pyabolism module.

### 2.2 The GC-Flux Algorithm

In the GC-Flux approach we manipulate the original optimization problem into a mathematically equivalent form, but one where every reaction is associated with a simple GPR string, which is either empty, a single gene or a single **AND** clause. This allows us to use expression data to impose constraints on our flux vector in a novel way, which better reflects the organisation of genes into functional complexes.

We first duplicate all reversible reactions, replacing each with one running in the forward direction and one with reactants and products exchanged. This results in a mathematically equivalent optimisation problem. While it is possible to constrain these pairs such that only one can run at a time, in this case we minimise the norm of the flux vector so that this additional constraint is not required. In this model, all elements of the flux vector are positive.

The second stage is to go through each reaction in turn and split the GPR string into individual viable gene complexes. As outlined in Sec. 2.1 this can be conveniently achieved by first converting the GPR string into a graph representation, before iterating over the directed graph to determine the appropriate list of gene-complexes. An example implementation of this can be found in the GC-Flux code of the Pyabolism python module. For example, if a reversible reaction had the GPR string **‘A or B’**, we would end up with four reactions, a forward and reverse reaction associated with each of the genes **A** and **B**.

We are now left with a model that has many more and different reactions than the original, but is mathematically equivalent to that found in Eq. 1. Applying either FBA or E-Flux to either model will give the same results as the unmodified model. However, this manipulation allows us to apply expression-related constraints in a principled way.

We still treat the RNA expression level as indicative of the protein's total flux-producing capability (not a model for the flux itself), but we now assume that this must place an upper limit on the rate across *all* reactions that rely on this gene for catalysis. Now that each gene only ever appears at most once in the GPR string for a given reaction, we can add the constraints separately to our linear program. We use the notation

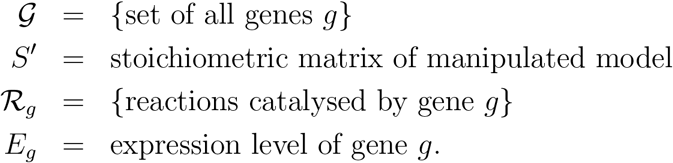

We are now in a position to define our GC-Flux problem in the following two-stage optimization problem. We first find *z*\*, the maximum achievable value of our chosen objective function.

#### Optimization 1.

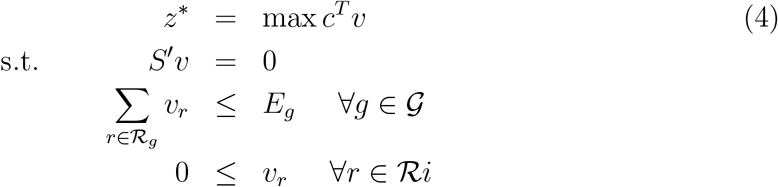

We then find υ*, the Flux vector of minimal magnitude that achieves this optimal flux.

#### Optimization 2.

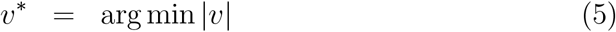

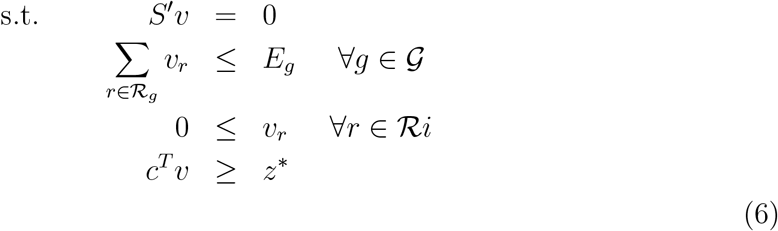

Figure 2 gives an intuition for this approach, considering each of the new reactions as pipes that expand to accommodate the flux they carry. Unconstrained, they can expand to accommodate any positive quantity of flux, but all those with GPR strings pass through hoops labelled with the relevant gene. The capacities of these hoops are set by the expression level of the respective gene, and there may often be multiple reactions passing through each hoop. The fluxes carried by reactions in the network are free to vary up the limits, but where multiple reactions are catalysed by the same gene, the total capacity must be divided between them. In this example, the capacity afforded by expression of gene ‘C’ must be shared between three different reactions.

**Figure 2:**
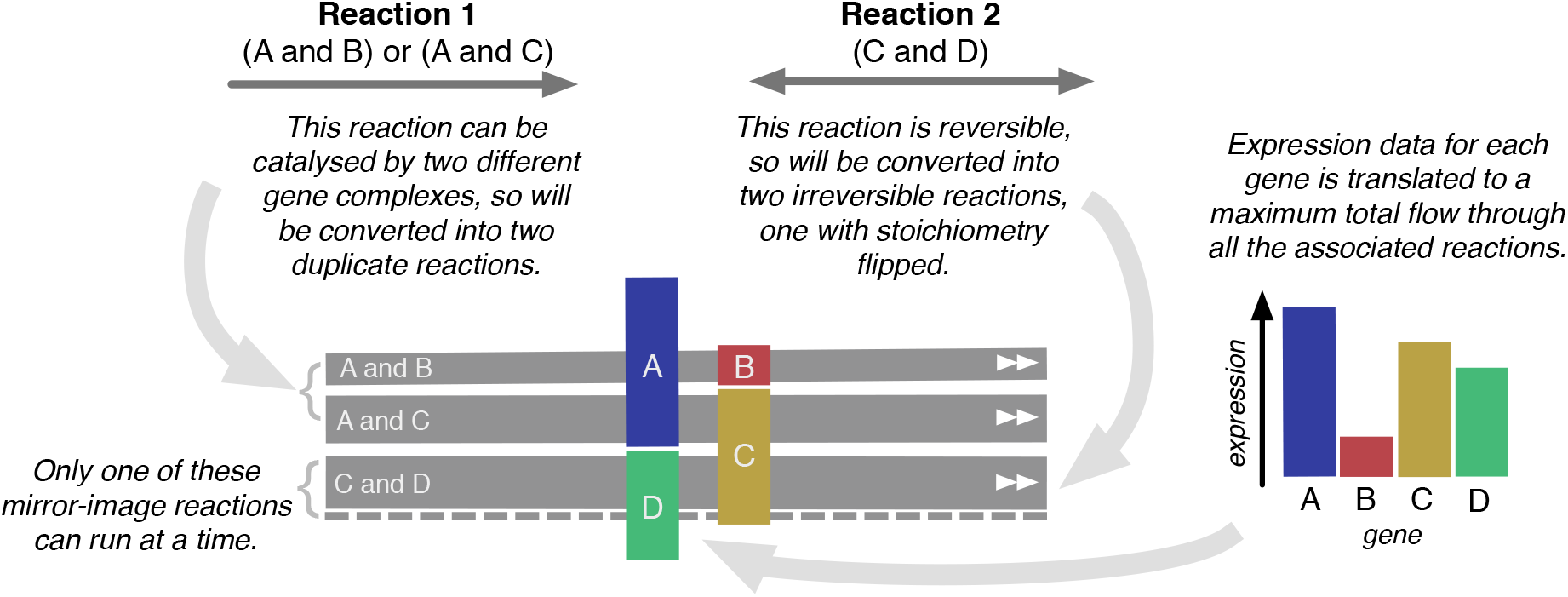
Illustration of how two original reactions relate to those in the converted model and the associated expression data. As indicated, the expression of each gene is directly proportional to the amount of flux permitted *in total* through all associated reactions. The ‘loops’ that constrain the fluxes are the same size as the corresponding bars on the chart showing expression levels. All reactions are uni-directional, and the dotted line indicates a reaction that is carrying no flux.

### 2.3 Verification with Experimental Data

To assess the performance of our algorithm, we use published experimental data for gene expression and metabolic fluxes in a variety of different conditions. This expression data provides input to the GC-Flux algorithm, and the correlation between predicted internal fluxes and those experimentally measured provides a metric for algorithm performance. We use data from two organisms across five different publications. For *Escherichia coli* (ecoli), we use transcriptomic data from Ishii et al. [24] and Holm et al. [22]. For the Ishii et al. study we use the microarray data which has wider coverage of genes, and covers five different knockout conditions. For *Saccharomyces cerevisiae* (yeast) we use data from [8] as well as the combined datasets from [46] and [26]. Henceforth we will refer to these datasets simply as Ishii, Holm, Celton and Rintalta, and we use the processed form provided by [29]. Kim et al.'s study gathers all the necessary data together as well as providing a mapping between measured reaction fluxes and reactions inside the models. Full details can be found in [29] and the associated supplementary material. As our ecoli model we use iJO1366 [42], and for yeast we use Yeast5 [21]. Details of all experimental conditions can be found in the original papers.

As a proof-of-concept demonstration we use a worst case scenario for such predictions, in which the only information we have on the growth conditions of the organism is the transcriptomic data. Such conditions could correspond to expression data from in-vivo growth, where bacterial growth occurs at an unknown rate on complex or uncharacterised media. This means no growth rate or carbon source specificity is known for the targeted cells, and hence the models we use in simulation do not have the corresponding constraints. For all algorithms we use the biomass vector included included in the curated models as the objective function.

We compare performance of GC-Flux to two existing algorithms, the afformentioned E-Flux2 [29] and **pFBA** [34]. The **pFBA** approach assumes that the organism will achieve the optimal objective, while minimising the resources used. Hence it performs standard FBA before minimising the flux through all reactions associated with enzymes. In order to allow comparison between the algorithms we choose to set an arbitrary uptake limit of 0.1 for all carbon sources, while every other exchange reaction with a non-zero limit is set to ±∞ as appropriate. The same non-expression constraints are used for all algorithms, meaning that we use a slightly adapted version of E-Flux2 in which uptake and ATP maintenance limits have been reintroduced.

The situation in which carbon source and growth rate is unknown removes a lot of information from the model, but there is still a large amount of organism-specific information tied up in the biomass vector. For commonly studied bacteria with well characterised metabolic models this special reaction has been carefully tuned, with a lot of experimental information and verification captured in its components and coefficients. To understand what predictive power this specific objective function brings to pFBA we also examine the capability of our algorithms to predict fluxes under an alternative objective. In line with the study of [49] we predict flux distributions from all three algorithms not only using biomass flux as an objective, but also maximsisation of ATP production.

We use multiple measures to assess the performance of the algorithms. Spearman correlation is the least stringent, assessing only the degree of monotonicity between silico and vitro fluxes. Pearson correlation indicates how well the data fits a linear model, and while the non-normality of the data means we cannot use the standard tests of statistical significance the measure nonetheless gives an indication of the how well the data fits a linear model. We also use the performance metric employed in [35], a normalised measure of the residuals between simulated and experimental fluxes. This takes the Euclidean norm of the residuals and normalises by the Euclidean norm of the experimental flux vector, allowing us to compare performance across the different experiments (which may have very different typical flux magnitudes).

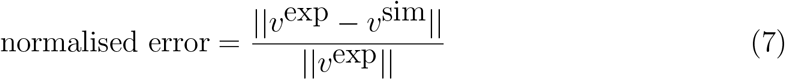

## 3 Results

Figure 3 shows the correlation achieved across all algorithms and objectives, with results displayed as violin plots showing not only the mean and degree of variation, but also how the results are distributed [55]. Performance is similar for all algorithms when biomass is employed as the objective, and this pattern is consistent across the three measures of performance. In contrast, when the objective is the production of ATP, we see a marked shift in performance between the algorithms. The pFBA approach shows a drop in performance, particularly for those metrics that are sensitive to the magnitude of error. In contrast, the integration of transcriptomic data to E-Flux2 and GC-Flux has made the predictions more robust to the choice of objective. This is not unexpected, given the relative lack of constraint when switching to a less sophisticated objective function, but it indicates that the predictive power of pFBA is at least partially due to the information contained in a well curated biomass vector.

**Figure 3:**
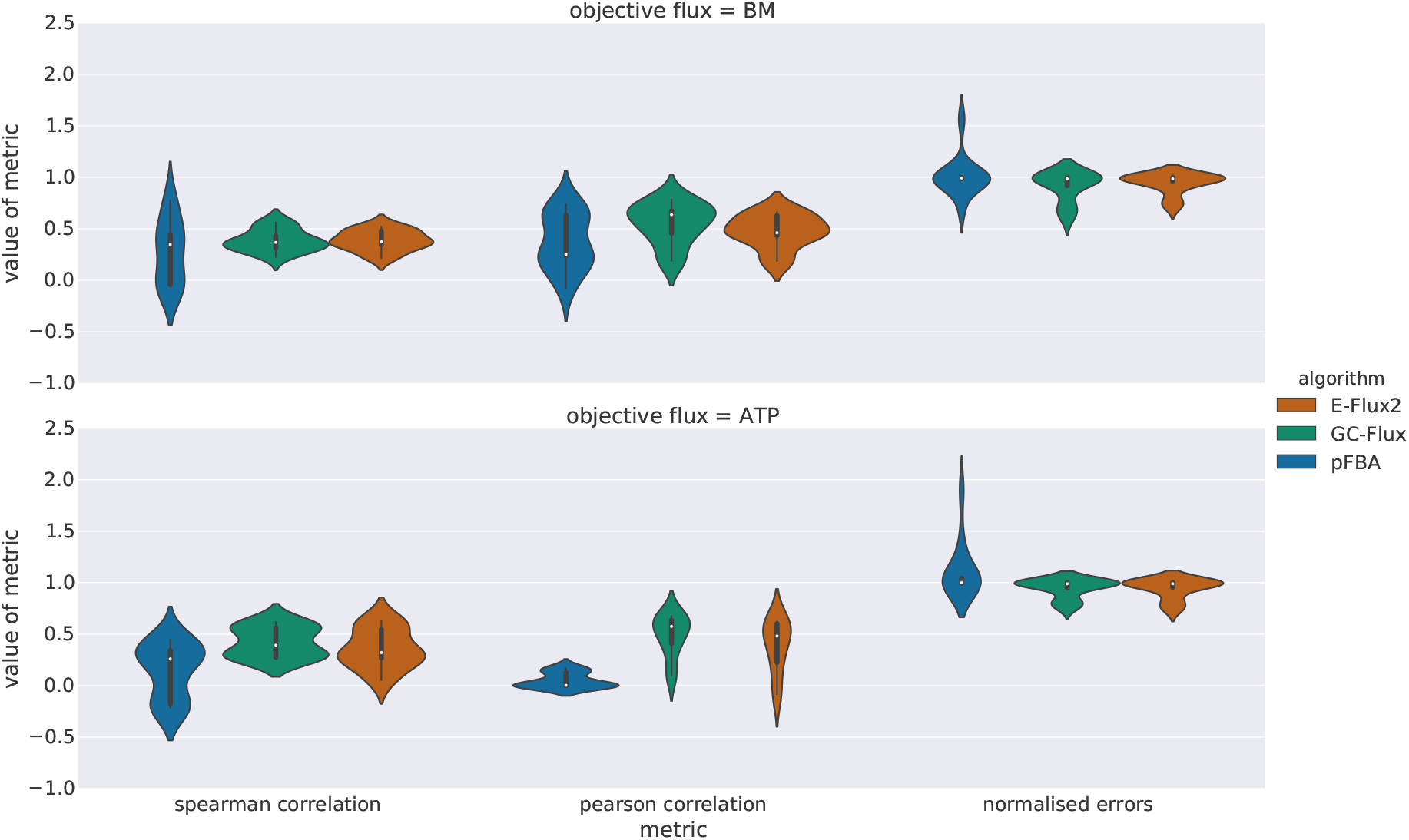
Results of flux vector predictions for the three algorithms with both biomass and ATP flux objectives. Both the Pearson correlation and normalised error metrics give the same qualitative pattern, in which performance is extremely similar when an appropriate biomass vector is known, but the expression-based techniques are much more robust when an alternative objective is employed.

### 3.1 Scaling Expression Data

Approaches that integrate transcriptomic data with constraint-based modelling have existing fixed constraints that impose, for example, ATP costs for cell maintenance [41], and where known, the uptake of a nutrient also provides a fixed constraint in defined units. The second type of constraint - on the internal reactions of the system, derived from transcriptomic measurements - is linked to the internal state of the organism, including expression and regulation of metabolic genes. Transcriptomic measurements do not naturally correspond to model units. Changing the relative values (via a conversion factor) of these two kinds of constraints changes the optimization problem in Eq. 4 and Eq. 5, and hence the predicted fluxes. To examine how this trade-off affects the quality of prediction, we rescaled the gene-centric constraints.

We performed the same set of simulations again, but with a range of linear scalings applied uniformly to the experimental expression values. For example, the constraints in the GC-Flux optimisation problem simply have a multiplicative factor added on the right side of the inequality, constant for all genes.

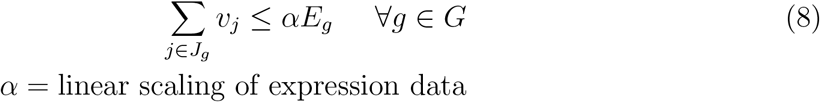

The bounds on E-Flux2 are similarly effected, and in our adapted version of the algorithm with absolute bounds on uptake and ATP maintenance, the scale-dependence is reintroduced.

Figure 4 shows the results of simulations for a wide range of scaling values, and displays a clear systematic variation in performance of both transcriptomic-based algorithms relative to the pFBA baseline. By definition the pFBA performance does not vary at all since it takes no account of the gene expression data, and in the limit of very large scaling the performance of both algorithms tends toward the pFBA value. At this limit the expression-derived bounds become so large as to be irrelevant as they are not providing any further constraint to the problem (the effective constraints are on the exchange reactions).

**Figure 4:**
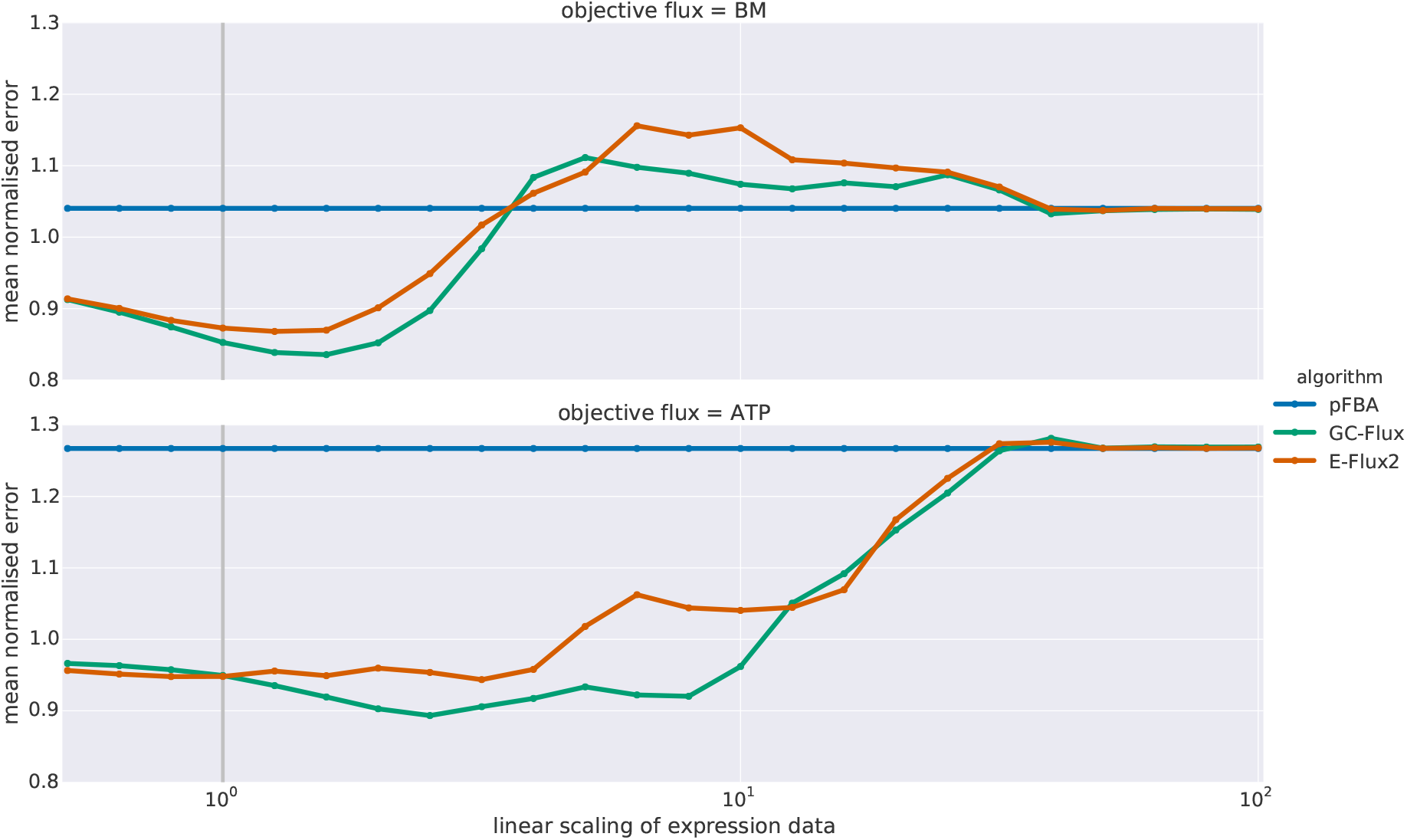
Variation in algorithm performance with uniform linear scaling of the transcriptomic data. As the magnitude of expression-derived constraints shifts relative to the inherent constraints we observe a non-monotonic shift in the performance of both E-Flux2 and GC-Flux. This clearly demonstrates that scaling of gene expression data, even when uniformly applied, can have a marked impact on the prediction of internal fluxes.

Figure 4 does not show an entirely straightforward relationship between performance and scaling value, but the ‘default’ scaling of 1.0 (used in Fig. 3 and previous studies [29, 35]) does not result in the best (peak) performance. Further, the peak performance achieved for **GC-Flux** is better in both cases than that of E-Flux2. As we vary the scaling we see fluctuations in performance, but given that applying the same scaling to all genes is somewhat blunt, this is not unexpected. The scaling that will ideally balance existing (exchange, ATP maintenance, thermodynamic) constraints with transcriptomic constraints will likely not be the same for all genes.

Transcriptomic data reflects the amount of mRNA present, and while this is overall likely to be indicative of metabolic activity and of flux capacity [10, 29, 38, 19, 18], the measured units are not directly comparable to that of flux constraints in FBA models. Scaling the expression data as we have done is a simple, global way of linking the two units. Our investigation gives an indication of what, on average, is the best scaling for the performance measures we have used, taken in the aggregate across all the genes/enzymes. Ideally, if individual kinetics were known, individual-level scalings for each gene, enzyme or complex could refine the constraints further. Our approach with a single global scaling is a first step, but can already improve flux prediction. In future it is possible that gene-or enzyme-specific scalings could even be learned from experimental observations combined with well-curated models, and that this could improve the use of transcriptomic data for future flux predictions.

## 4 Conclusions

We have introduced a novel formulation combining experimental gene expression data with constraint-based metabolic modelling, in which viable gene complexes are considered as the fundamental unit. This relegation of the status of reactions allows more realistic use of expression data, acknowledging that the individual proteins present in cells cannot be simultaneously employed in multiple reactions.

Using expression data to set limits on reaction rates is fraught with difficulty, and such data can only really be seen as indicative rather than truly quantitative. As discussed in [5], RNA expression data is far from perfectly correlated with intracellular protein concentration, but despite the post-translational factors such as protein half-lives and sRNA regulation contributing to discrepancies between mRNA and protein abundance [36], the literature indicates a degree of positive correlation, both in yeast [19, 18] and prokaryotic organisms [38, 39]. In both E-Flux2 and GC-Flux the limits are set as *maximum* rates rather than *realised* flux rates, leaving scope for system-level effects to prevent the full utilisation of reactions’ capacities. Despite the limitations, treating such experimental data as quantitatively predictive can give useful and verifiable results.

We have worked in the setting in which the carbon source and growth rate are not known, as well as examining performance in cases when the biomass objective function is absent. While this information would typically be available in many culture-based experiments, there is growing interest in using constraint-based metabolic modelling approaches to understand the function and dynamics of communities of microbes described with metagenomics [33, 54]. As metagenomic tools cannot preserve linkage information, it is challenging to reconstruct which species are present in a sample [13]; many species present are unknown and/or cannot be cultured. It is currently prohibitive to define high-quality species-specific metabolic models to capture the metabolic capabilities of the microbial communities that are characterised with metagenomics. Consequently, moving to a gene-centric approach, as we have done here, has a natural appeal for applying metabolic modelling tools to metagenomic datasets.

The GC-Flux approach also opens up the possibility of tuning expression-derived constraints individually, learning the appropriate coefficients that would translate individual mRNA measurements into more accurate constraints on the maximum reaction rates. Each reaction r ∈ *R* would have an associated constant *k_r_* ∈ *K,* which determines how fast a particular value of the expression of the gene-complex will permit the reaction to run.

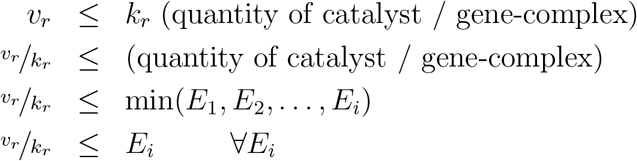

The constant *k_r_* is a property of the reaction, and incorporating this into the framework of optimisation problems 1 & 2, we define new optimisation problems that take these ‘kinetic constants’ into account.

### Optimization 3.

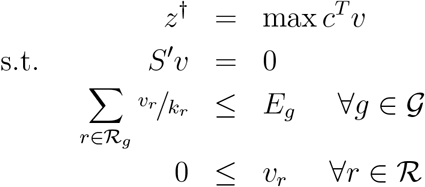

### Optimization 4.

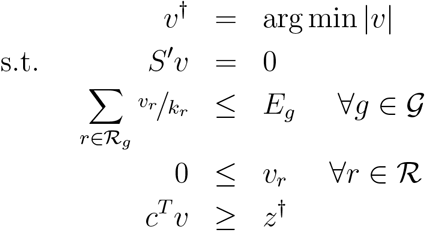

We are left with the task of finding a suitable set of constants *K* such that in solving Optimisations 3 & 4 we obtain a flux vector with the best possible correlation with experimental results.

An approach to solving this problem could provide a step toward high-throughput determination of kinetic parameters in metabolic networks, permitting improved dynamic modelling (e.g. with ordinary differential equations). This multi-level optimisation is not easily tackled, and may require the introduction of a suitable approximate form, but it represents an interesting avenue for future research.

GC-Flux represents a conceptual advance in constraint-based modelling, giving a principled basis for linking expression data with maximum Flux capacities. We have demonstrated the algorithm on experimental data, showing improved predictive power when multiple representation of genes in models is taken into account. A shift to focusing on gene complexes as the fundamental unit of constraint-based metabolic modelling is both theoretically appropriate and practical.

## Acknowledgement

### Funding

This work was supported by the Biotechnology and Biological Sciences Research Council of the United Kingdom (NF and CC: grant BB/I00713X/2). This paper was substantially improved by the detailed comments of anonymous reviewers.

